# Polysome Profiling Method for Low-Input Human Postmortem Brain

**DOI:** 10.64898/2026.05.28.726378

**Authors:** Vandana Sharma, Anuj Choudhary, Mrunali Dinesh Dhokne, Barbara Gisabella, Harry Pantazopoulos, Rammohan Shukla

**Author notes:** These authors contributed equally for this position (Anuj Choudhary, Mrunali Dinesh Dhokne). **Corresponding author’s email address** (Vandana Sharma), (Rammohan Shukla).

## Abstract

Polysome profiling is a powerful technique used to analyze the association of mRNA with ribosomes, providing insights into the translational status of a cell. It relies on the separation of ribosome-bound mRNAs through sucrose density gradient centrifugation, where the number of ribosomes on an mRNA correlates with its sedimentation rate. While numerous studies have successfully applied this method to cell line and mouse tissue, application to the human postmortem brain remains scarce due to challenges related to sample quality and low concentration of recoverable material. To overcome these challenges, we:

1. Implemented a protocol specifically optimized for low-concentration human post-mortem brain tissue.
2. Implemented a gradient-maker-free method to manually prepare sucrose gradients with tunable sensitivity for low-input samples.
3. Adapted the human brain tissue protocol for neuronal cell lines and mouse brain with minimal modification.

## Background

Regulation of mRNA translation plays a central role in gene expression, influencing critical biological processes such as development, synaptic plasticity, cellular stress responses, and disease pathogenesis [1]. Under conditions such as nutrient deprivation, oxidative stress and various disease states, including cancer and neurodegenerative disorders, translation is frequently dysregulated.

Ribosomes, the translational machinery, embody core principles of translational regulation. They exist as distinct ribosomal assemblies along the translational pathway, and the distribution of these assemblies reflects the translational landscape of a cell [2]. For example, 40S and 60S subunits mark pre-initiation or termination phases, 80S monosomes typically represent translation initiation, disomes and trisomes can indicate ribosome pausing or stalling, and polysomes reflect active elongation.

Polysome profiling by separating different ribosomal assemblies based on their sedimentation through ultracentrifugation in a sucrose density gradient provides a quantitative snapshot of this translational landscape within a given sample. Peaks corresponding to ribosomal subunits, monosomes, and polysomes can be resolved, and the ratio of monosome to polysome can be used as a proxy for translational efficiency, enabling quantification of global shifts in translation under physiological or pathological conditions.

Existing polysome profiling protocols, widely used in cultured cells, often require optimization when applied to complex tissues [3,4]. Human postmortem brain samples present distinct constraints, including limited input, sample heterogeneity and variable RNA integrity, which together restrict the broader application of polysome profiling in this context. As a result, studies using postmortem tissue have predominantly focused on RNA or protein measurements, leaving translational regulation largely unexplored. Adapting polysome profiling for human postmortem samples provides a critical opportunity to access this missing layer of gene expression control, particularly in the context of neurological and psychiatric disorders.

Here, we present a streamlined and adaptable protocol for polysome profiling of human postmortem brain tissue. The method enables reproducible separation of translational fractions using manually prepared sucrose gradients, optimized for low-input and implementable without specialized gradient making equipment, and is also applicable to cultured cells and mouse brain.

## Method details

### Step 1: Sucrose gradient preparation

#### Materials

- Gradient buffer (Table S1)
- Sucrose (Sigma Aldrich, Cat#: S0389, molecular biology grade; dilutions: Table S2)
- DEPC (Sigma Aldrich, Cat#: 40718; DEPC water [Table S3])
- Minisart Syringe Filters (Fisher Scientific, Cat#: 14555264)
- Polyclear™ Open Top Ultracentrifuge Tubes SW41 9/16″ X 3 1/2″ (Seton Scientific, Cat#: 7030)
- Tube stand (snug-fitting; VWR, Cat#: 89215-718)
- Magnetic stirrer
- Dry ice or −80°C freezer

Standard gradient preparation often relies on automated gradient makers, which are widely used to generate continuous sucrose gradient across the full length of the tube (Figure 1A). However, these systems require dedicated instrumentation not readily available in all laboratories. To provide an accessible and equipment-independent alternative, we implemented a “step-gradient” tailored to sample input. In this approach, sucrose layers of defined concentrations are added in adjustable volumes (Figure 1B and C) allowing controlled redistribution of gradient density within a defined region of the tube. Upon equilibration, these layers form a continuous gradient comparable to that produced by the gradient makers. Because individual layer volumes can be modified without instrument recalibration or reprogramming, this strategy offers practical flexibility for low-input optimization while maintaining reproducibility.

**Figure 1.**
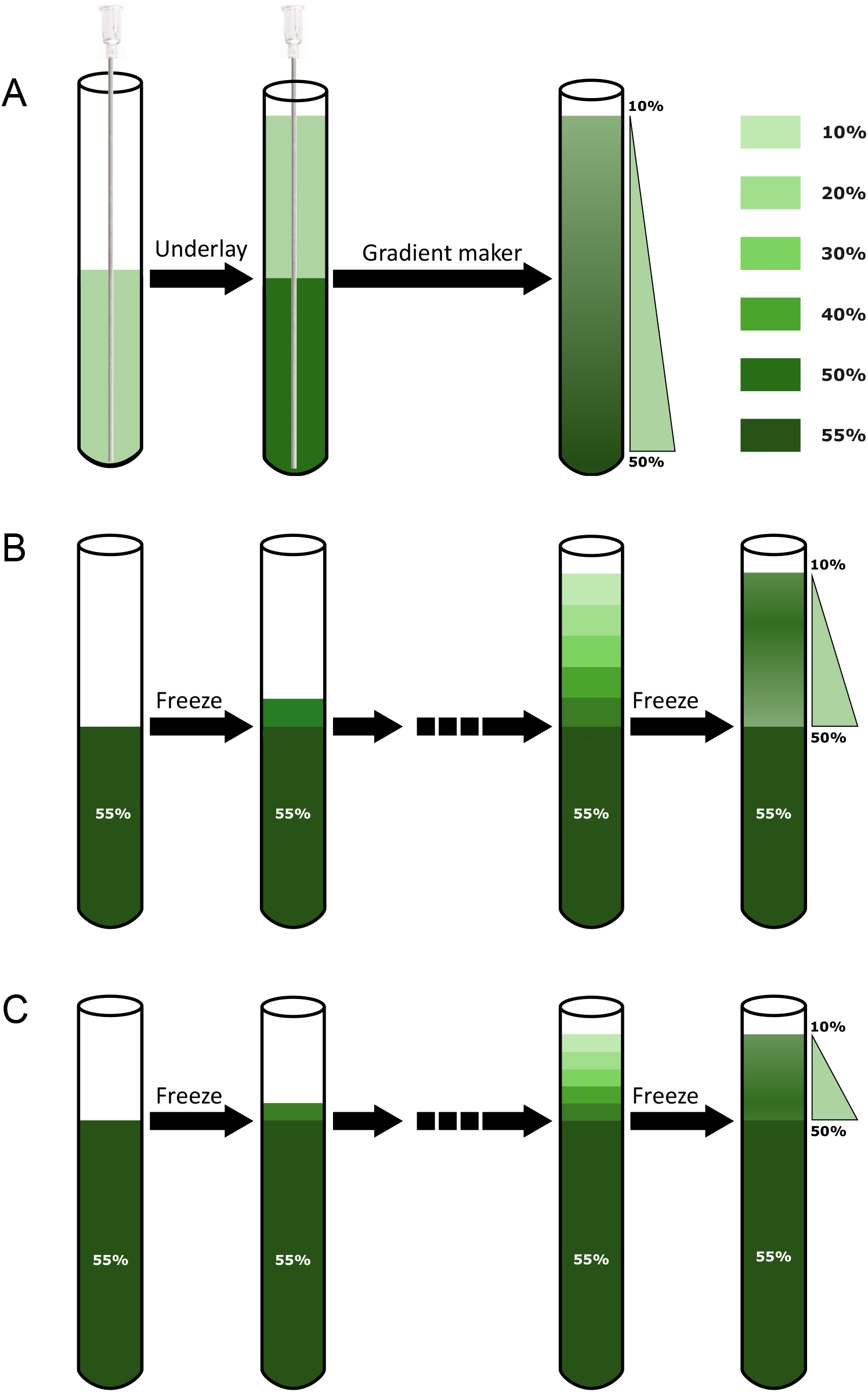
Comparison of automated and manual step-gradient preparation strategies: **(A)** Automatic gradient: Continuous sucrose gradients were prepared by first adding a 10% sucrose solution, followed by underlaying 50% sucrose using a Hamilton needle. A gradient maker was then used to generate a continuous gradient. **(B)** Manual gradient: Step sucrose gradients were prepared by sequentially adding sucrose solutions of decreasing concentration into ultracentrifuge tubes (in order: 8 mL of 55%, followed by 0.8 mL each of 50%, 40%, 30%, 20%, and 10%). After the addition of each layer, tubes were placed at ™80°C to freeze before adding the next layer, and were finally stored at ™80°C. Prior to use, gradients were equilibrated overnight at 4°C. **(C)** Manual compressed gradient: Gradients were prepared as in (B), except that 10.5 mL of 55% sucrose and 0.3 mL each of 50–10% sucrose were added.

The gradient is prepared by sequentially freezing sucrose layers of varying concentrations using dry ice or a −80°C freezer. We begin with filling 55% sucrose, allow it to freeze, and then sequentially loading and freezing layers from 50% to 10% sucrose (Figure 1B). Since polysomes do not migrate beyond 55% sucrose, this layer serves as a cushion to support and adjust the upper steps. The cushion volume (55% sucrose) can be increased, and the step volumes (50–10%) can be decreased, to accommodate lower input samples (Figure 1C). Gradients can be prepared in batches and stored at −80°C for up to one month.

1. Prepare 55% sucrose in gradient buffer (Table S1). Dissolve the sucrose using a magnetic stirrer for ~20 minutes. Do not apply heat to dissolve sucrose, as heating can alter the pH of the buffer. Approximately 70 mL of 55% sucrose is sufficient to layer six ultracentrifuge tubes and prepare all required dilutions (Table S2).
2. Filter the prepared 55% sucrose solution using a 0.22 or 0.45 µm filter either with a syringe filter or a vacuum filtration unit.
3. Prepare 10–50% sucrose solutions by diluting the 55% sucrose according to the recipe in Table S2. 5 mL and 2 mL of each dilution is sufficient for six tubes of 0.8 mL and 0.3 mL step gradient, respectively.
4. Rinse ultracentrifuge tubes with DEPC-treated water (Table S3) before use.
5. Place tubes in a snug-fitting tube stand to prevent movement during layering (Figure S1A).
6. Prepare the step gradient by sequentially adding sucrose solutions from 55% to 10%, in decreasing order of concentration (Figure 1B).
7. After adding each sucrose layer, freeze the tube using dry ice (or a −80°C freezer if dry ice is unavailable) before adding the next layer.
8. Repeat the “layer-and-freeze” process (step 6 and 7) until all gradient steps are added.
9. Seal the prepared gradients with paraffin film and store at −80°C.
10. Equilibrate gradients at 4°C overnight (≥ 12 hours) prior to use.

### Step 2: Sample preparation

#### Materials

- Human postmortem brain tissue
- Mortar and pestle
- Liquid Nitrogen
- Lysis buffer (Table S4)
- 23-gauge needle
- 1 mL syringe
- Refrigerated microcentrifuge
- Microcentrifuge tubes (1.5 mL)
- KIMBLE Dounce tissue grinder set (Sigma Aldrich, Cat#: D8938-1SET), including loose-(A) and tight-pestle (B)
- Nanodrop One Spectrophotometer (Fisher Scientific, Cat#: 13400525S24)

The use of human postmortem brain tissue for polysome profiling requires careful consideration of RNA integrity, which can vary widely across samples. Based on prior observations and published report [5], brain sections were first pulverized under liquid nitrogen prior to lysis. Pulverization enhances extraction efficiency and provides more accurate control over the amount of tissue used for downstream processing. We selected tissue with RNA Integrity Number (RIN) values of 6.5 or higher (Figure S1B), a range generally accepted for postmortem brain analysis [6,7].

All steps, from tissue thawing to centrifugation, should be carried out on ice to preserve ribosome integrity and prevent RNA degradation. For efficient lysis and good polysome peak resolution, sufficient lysis buffer (in the ratio of 1:5) is critical. To minimize RNase contamination, all plasticware and glassware should be rinsed with DEPC-treated water. Lastly, as an end point of this step, sample input should be measured using RNA concentration. Unlike other protocols that normalize sample input based on optical density (OD) measurements [3], RNA concentration serves as a more reliable guide for sample loading onto sucrose gradients.

1. Pulverize a small piece of brain tissue using a mortar and pestle under liquid nitrogen. Transfer the pulverized tissue to a pre-chilled microcentrifuge tube and weigh the required amount for downstream processing.
2. Transfer the required amount of pulverized tissue to a pre-chilled glass Dounce homogenizer kept on ice.
3. Add lysis buffer (Table S4), using at least 50 µL per 10 mg of pulverized tissue, and incubate on ice for 5 minutes.
4. Using the loose pestle (A), slowly and steadily homogenize the pulverized tissue in the pre-chilled Dounce homogenizer kept on ice until it is fully resuspended in lysis buffer and no visible clumps remain (approximately 5 minutes).
5. Next, use the tight pestle (B) to homogenize again for an additional 5 minutes (slowly).
6. Transfer the supernatant to a chilled 1.5 mL tube.
7. Using a 23-gauge needle, triturate the lysate 10 times, then incubate on ice for 10 minutes.
8. Centrifuge the lysate at 16,000 x g for 6 minutes at 4°C. Transfer the supernatant to a chilled 1.5 mL tube.
9. Repeat step 8 to further remove insoluble material.
10. Measure the RNA concentration in the tissue lysate using a UV-Vis Nanodrop spectrophotometer.

For best results, use the lysate on the same day. However, if necessary, it can be stored at −80°C for later use, provided it is frozen immediately after preparation.

### Step 3: Loading and balancing the gradient and running the ultracentrifuge

#### Materials

- Optima L-80 XP Preparative Floor Ultracentrifuge (Beckman Coulter)
- SW 41 Ti Swinging-Bucket Rotor (Beckman Coulter)
- A 2-decimal place balance (Sartorius Entris II Essential Precision Balance, Cat#: BCE2202I-1S)
- Super Lube-21030 Synthetic Multi-Purpose Grease

To preserve the gradient integrity, polysome profiling is performed using a swinging bucket rotor. To maintain temperature stability and streamline the workflow, keep the ultracentrifuge rotor and buckets at 4°C when not in use. Proper balancing of the ultracentrifuge tubes containing the prepared step gradient (from step 1) is critical for safe and accurate operation. While balancing, ultracentrifuge tubes paired opposite to each other in the rotor must be adjusted with sample lysate or lysis buffer to ensure equal weight. Balancing is performed with the ultracentrifuge tubes placed inside their respective buckets. Each tube should be filled to at least 95% of its total volume to prevent collapse under high-speed conditions (Figure S1A). For consistency, it is advisable to complete the balancing procedure briskly to minimize temperature fluctuations.

To minimize the time between centrifugation and fraction collection, start and prime the fractionator (Step 4) as soon as the run is completed, and while the ultracentrifuge is decelerating (typically ~20 minutes). After removal from the rotor, place the buckets in the provided bucket rack to avoid disturbing the gradient. Because fractionation is performed sequentially and each tube requires processing time, keep the bucket rack in an ice box containing ice-cold water covering approximately three-quarters of the bucket height to maintain temperature stability until all tubes are processed.

1. Start the vacuum system and pre-cool the ultracentrifuge to 4°C. This may take approximately 20 minutes.
2. Remove the rotor buckets from the refrigerator and gently insert each ultracentrifuge tube containing the sucrose gradient (from step 1).
3. Using an analytical balance, carefully balance each rotor bucket with the ultracentrifuge tube placed inside it, ensuring that the corresponding cap (pre-greased if required) is included during weighing.
4. Load the sample lysate onto the top of each sucrose gradient so that each tube is filled to within 2–3 mm from the top. Approximately 125 µL of sample corresponds to 1 mm of height.
5. Balance the opposing tube as needed using lysis buffer.
6. Remove the rotor from the refrigerator and wipe off any condensation.
7. Load the balanced buckets into the rotor, ensuring they are hooked properly.
8. Stop the ultracentrifuge vacuum system, then carefully place the rotor inside.
9. Set acceleration to maximum and deceleration to level 5. *NOTE: setting the deceleration level to 5 is very important to maintain the gradient when the run is finishing*.
10. Run ultracentrifugation at 36,000 rpm for 2 hours at 4°C.
11. After the run, carefully unload the rotor and place the buckets in ice water along with the rotor rack.
12. When ready for fractionation, gently remove the tubes from the buckets using tweezers, and proceed to step 4.

### Step 4 Gradient fractionation

#### Materials

- Piston Gradient Fractionator (BioComp Instruments, Cat#: 153)
- Gilson Fraction FC-203B Collector (BioComp Instruments, Cat#: 151-203)
- Triax Flow Cell (BioComp Instruments, Cat#: FC-3-2UV-VIS)
- 96 well plate holder (BioComp Instruments, Cat#: 151-203-96)

Before loading your samples, it is recommended to run a water-filled tube to ensure the system is functioning properly and to avoid wasting valuable samples. Depending on the downstream application, fractions should be collected in volumes that allow clear separation of the desired peaks—such as 80S, light-, and heavy-polysomes. Collecting few fractions with large volumes can cause adjacent peaks to overlap, reducing separation and resolution.

The Biocomp fractionator, unlike models that pierce the tube, operates on the principle of positive displacement, allowing ultracentrifuge tubes to be reused—typically for up to six runs. To prevent degradation, immediately store collected fractions at 4°C for short term storage and −80°C for long-term storage.

For gradient prepared using gradient maker, fractionation and instrument setup followed the manufacturer’s operating guidelines and instructional resources [BioComp Instruments Ltd.; available at https://www.youtube.com/@biocompinstr0075mentsltd.1869]. For manually prepared step gradients, the same fractionation workflow can be followed; however, it is recommended to define the “Zone of fractionation” in the “SCAN” tab of the software to restrict data acquisition to the gradient region. (Figure 2A). This setting allows the scanner to process only the resolved gradient layers while excluding the high-density cushion, thereby reducing overall fractionation time. A concise overview of the workflow used in this study is provided below.

**Figure 2.**
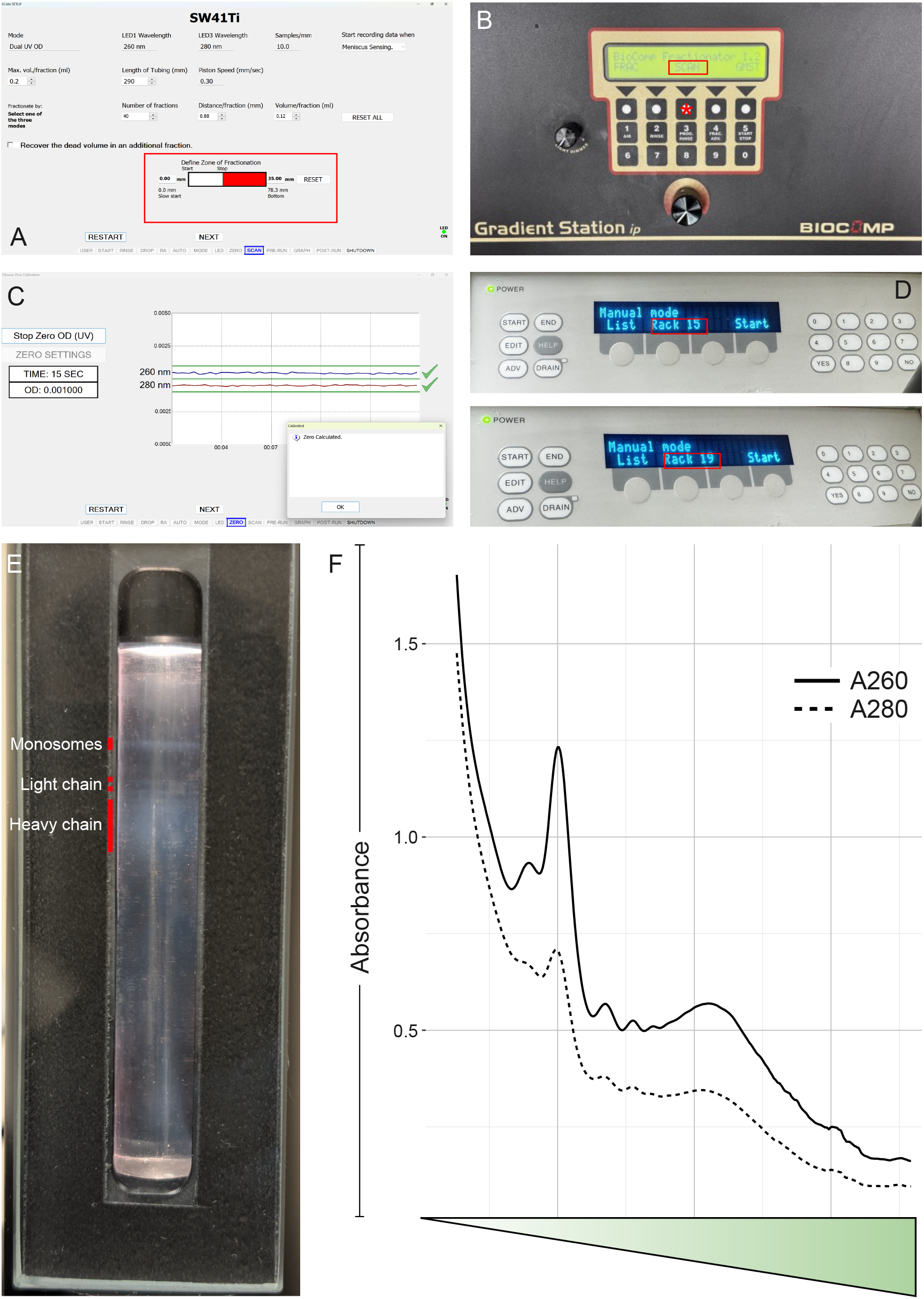
Fractionation workflow using the BioComp piston gradient fractionator. **(A)** Defining the “Zone of fractionation” to restrict scanning to the gradient region. **(B)** Initiating system communication by pressing “SCAN.” **(C)** Stabilizing the UV baseline at zero OD. **(D)** Selecting the appropriate collection rack (15 for 96-well plates; 19 for microcentrifuge tubes). **(E)** Visualization of ribosomal assemblies prior to fractionation. **(F)** Representative A260 (RNA, solid) and A280 (Protein, dashed) absorbance profiles obtained during fractionation.

1. Turn on fractionator and computer. Press “SCAN” on the gradient station control panel (Figure 2B) to connect the fractionator to the computer.
2. Launch the Triax™ FlowCell software and follow the on-screen instructions.
3. Rinse the system thoroughly with DEPC-treated water to remove residual material and stabilize the UV detector baseline at zero absorbance (zero OD; Figure 2C) before sample fractionation. *NOTE: If you notice spikes during zeroing or in the polysome profile, rinse with 20 mL of 1 M NaOH followed by 50 mL of DEPC-treated water before proceeding*.
4. Prepare 1.5 mL tubes or a 96 well plate for collection and place them in the respective rack on the fraction collector. Select rack number 15 for the 96 well plate and rack number 19 for microcentrifuge tubes (Figure 2D).
5. Gently remove the centrifuge tube from the ultracentrifuge bucket stored on ice, taking care not to disturb the gradient.
6. Attach the SW41 tube holder cap onto the centrifuge tube and securely place the capped tube into the holder.
7. Insert the loaded tube holder into the fractionator and rotate clockwise to secure it. At this stage, for the manually prepared step gradient, turning on the piston tip light allows visualization of distinct monosome rings and diffuse polysome regions within the tube when sufficient material is present (Figure 2E). *NOTE: Avoid keeping the piston tip light on for too long to prevent RNA degradation from heat*.
8. Begin fraction collection by clicking “START SCAN” in the software, which collects each fraction from top to bottom and records UV absorbance at 260 nm and 280 nm. Absorbance at 260 nm reflects RNA content and is the primary readout for polysome profiling. The 280 nm signal corresponds mainly to protein and typically mirrors the 260 nm peak pattern in ribosomal fractions, serving as an internal confirmation that the observed peaks represent ribonucleoprotein complexes rather than background noise (Figure 2F). *NOTE: Clean and dry the piston tip between runs to prevent sticking during scanning*.
9. Immediately place collected fractions on ice and store at −80°C until further use.

### Adaptation for cell-lines and mouse brain tissue

For mouse brain tissue, follow the same steps as outlined for human brain tissue without any modification. When working with primary cells or immortalized cell lines for polysome profiling, ensure that cells are in the logarithmic growth phase (60–70% confluency) before the experiment. Higher confluency can lead to reduced polysome peaks due to decreased proliferation and lower global translation activity. Unlike brain tissue, cultured cells should be treated with cycloheximide (100 µg/mL) for 5–15 minutes prior to harvesting to stabilize ribosomes on mRNA. The optimal cycloheximide treatment time should be standardized for each cell type. While it is best to use freshly harvested cells for robust polysome profiles, cells may also be flash-frozen and stored at – 80°C until use. Notably, sodium deoxycholate can be excluded from the lysis buffer, as cultured cells are easily lysed with non-ionic detergents.

## Data analysis

After obtaining the polysome profile on the Biocomp Gradient Station, the UV absorbance trace can be exported as a .csv file via the TRIAX software, which comes with the Biocomp Gradient Station. While the software allows for overlaying profiles, merging profiles from up to four runs, and downloading the aligned data for use in Excel, it cannot perform downstream quantitative analysis. Specifically, it cannot compute the area under the curve (AUC) or calculate the monosome-to-polysome (M/P) ratio. Both metrics are essential for comparing peaks and interpreting differences between translational states. These calculations can be performed using quAPPro, an open-source R Shiny platform that can calculate AUC values, determine the M/P ratio, and conduct statistical tests using the full raw data downloaded from the TRIAX software [8]. The documentation and tutorial for quAPPro is publicly available online via its GitHub repository.

### Method validation

To validate the applicability of the step-gradient approach in human postmortem brain tissue, polysome profiles were generated using automated gradients (Figure 3A and D) and two manual configurations differing in step volume: 0.8 mL standard (Figure 3B and E) and 0.3 mL compressed (Figure 3C and F). At an input of 300µg RNA, the automated gradient resolved a distinct monosome peak but showed comparatively limited separation of light and heavier polysome fractions. In contrast, both manual step gradients, particularly the compressed 0.3 mL configuration, produced clearer delineation of monosome, light polysome (disome/trisome), and heavier polysome regions. Importantly, this pattern of segregation was maintained at lower input (50µg RNA), where the manual gradients continued to resolve distinct ribosomal assemblies.

**Figure 3.**
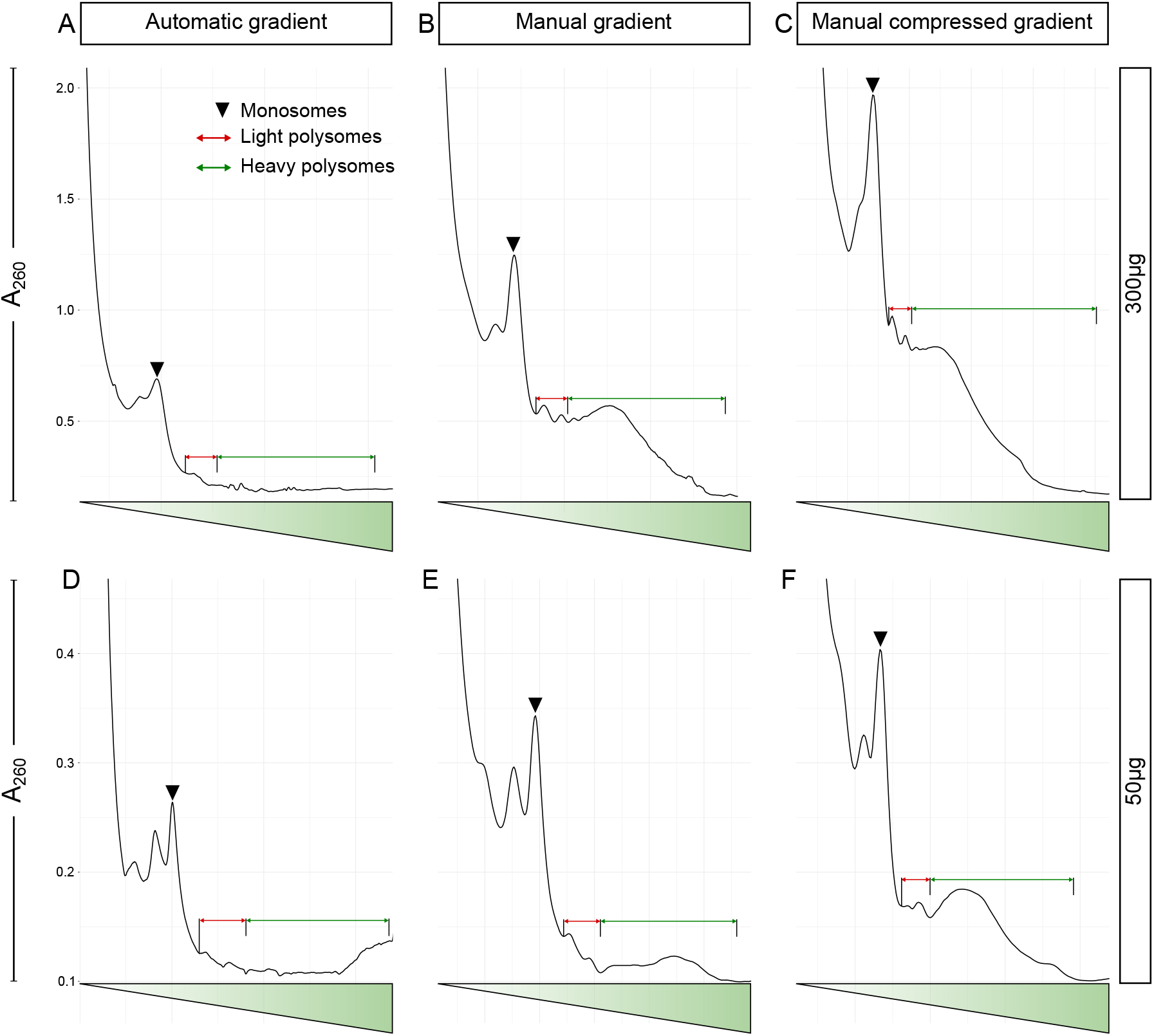
Validation of step-gradient performance in human post-mortem brain samples. Representative A260 polysome profiles generated using automatic, manual, and manually compressed gradients at 300µg **(A–C)** and 50µg **(D–F)** RNA input. Note that manual and compressed step gradients show improved delineation of monosome, light polysome, and heavier polysome regions across input amounts compared with the automatic gradient.

### Limitations

Although steps smaller than 0.3 mL could allow for use of even lower inputs, they reduce the resolution of peaks representing disomes, trisomes, tetrasomes, and higher order polysomes, and complicate fraction collection. We have not tested the protocol in samples with RNA Integrity Number (RIN) values below six. Based on the known impact of RNA degradation on ribosome loading, we believe these samples would display a disproportionately high M/P ratio. This would likely reflect reduced polysome abundance, as longer mRNAs are more suspectable to degradation, along with increased monosome levels due to fragmented mRNAs preferentially accommodating single ribosomes.

## Ethics statements

No live human participants or animals were involved in this study. Human postmortem brain tissue was obtained from the University of Mississippi Medical Center (UMMC) Brain Bank, in accordance with institutional and national ethical guidelines, with all necessary approvals in place.

## CRediT author statement

Conceptualization: RS and VS; Methodology: VS; Resources: HP and BG; Investigation: VS, MDD and AC; Data Curation, VS and RS; Writing – Original Draft Preparation, VS and RS; Supervision, RS and VS; Project Administration, RS; Funding Acquisition, RS.

## Acknowledgments

This work is supported by Institutional Development Award from the National Institute of General Medical Sciences of the National Institutes of Health under Grant # 2P20GM103432 and 2P20GM121310 to RS.

## Declaration of interests

The authors declare that they have no known competing financial interests or personal relationships that could have appeared to influence the work reported in this paper.

## Supplementary material *and/or* additional information

Additional figures and tables supporting the manuscript are provided as supplementary information.

## Notes

### Competing Interest Statement

The authors have declared no competing interest.

