## Supplementary material for "Polysome Profiling Method for Low-Input Human Postmortem Brain": Sharma_Protocol_Supplementary-Info.pdf

**Supplementary Information**

**Buffer Formulations and Sucrose Gradient Preparation Details**

**Table S1: Gradient Buffer (pH 7.4)**

| Reagent | Final concentration | Amount |
|---|---|---|
| Tris-HCl pH 7.4 (1 M) | 20 mM | 1.4 mL |
| $MgCl_2$ (1 M) | 5 mM | 0.35 mL |
| NaCl (4 M) | 150 mM | 2.62 mL |
| Glycerol | 8 % | 5.6 mL |
| *DTT (500 mM) | 1 mM | 0.14 mL |
| *Cycloheximide (100 mg/ml) | 0.1 mg/ml | 0.7 mL |
| DEPC $H_2O$ | n/a | 59.19 mL |
| **Total** | | **70 mL** |

**Table S2: DEPC-treated water**

| Reagent | Final concentration | Amount |
|---|---|---|
| DEPC | 0.1 % | 1 mL |
| dd$H_2O$ | n/a | 999 mL |
| **Total** | | **1000 mL** |

*NOTE: DEPC does not dissolve immediately in water; stir (preferably overnight) until the globules disappear. Autoclave to degrade DEPC and allow the solution to cool to room temperature before use.*

**Table S3: Sucrose Gradient Formulations (10–50%) Prepared from 55% Stock**

| Sucrose | Amount of 55% sucrose | Amount of gradient buffer |
|---|---|---|
| 50% | 4.54 mL | 0.46 mL |
| 40% | 3.63 mL | 1.37 mL |
| 30% | 2.72 mL | 2.28 mL |
| 20% | 1.81 mL | 3.19 mL |
| 10% | 0.90 mL | 4.10 mL |

*NOTE: 5 ml of 10-50% sucrose buffers are sufficient to prepare 6 tubes (SW41Ti).*

**Table S4: Lysis Buffer (pH 7.4)**

| Reagent | Final concentration | Amount |
|---|---|---|
| Tris-HCl (1 M) pH 7.4 | 20 mM | 20 µL |
| $MgCl_2$ (1 M) | 5 mM | 5 µL |
| NaCl (4 M) | 150 mM | 37.5 µL |
| Glycerol | 8% | 80 µL |
| Triton-X (10%) | 1% | 100 µL |
| #Sodium deoxycholate (10%) | 1% | 100 µL |
| *DTT (500 mM) | 1 mM | 2 µL |
| *Cycloheximide (100 mg/mL) | 100 µg/mL | 1 µL |
| *PMSF (200 mM) | 1 mM | 5 µL |
| *Protease inhibitor EDTA free (10X) | 1 X | 100 µL |
| *Turbo DNAse (2U/µL) | 24 U/mL | 12 µL |
| *RNase Inhibitor (40U/µL) | 100 U/mL | 2.5 µL |
| DEPC $H_2O$ | n/a | 632.5 µL |
| **Total** | | **1 mL** |

*# Used only for brain tissue; *Add fresh, just before use*

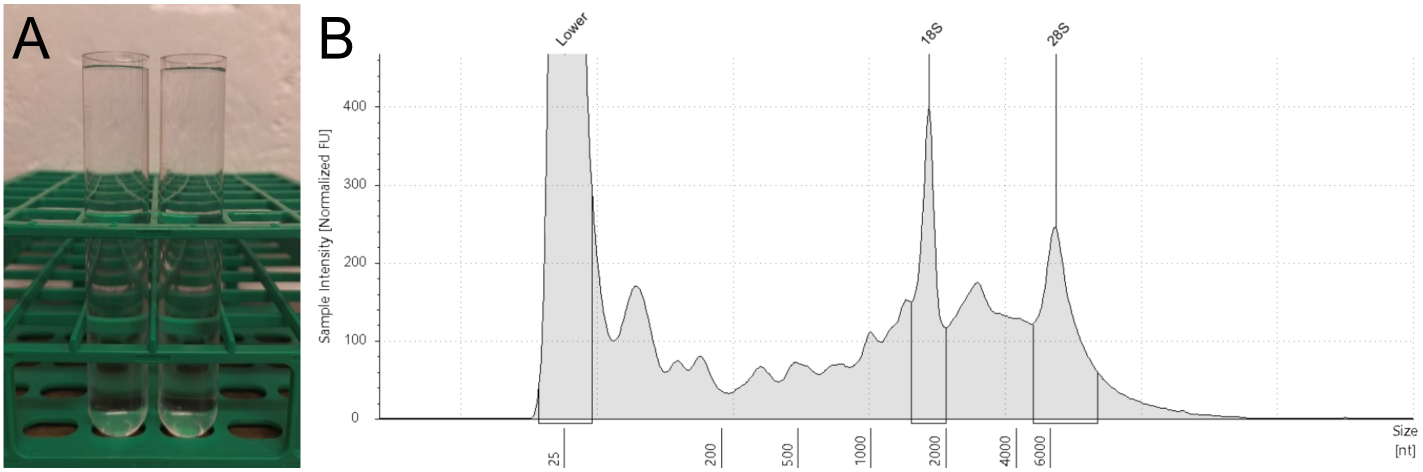


Sample Table

| Well | RINe | 28S/18S (Area) | Conc. [ng/µl] | Sample Description | Alert | Observations |
|---|---|---|---|---|---|---|
| H8 | 6.7 | 1.2 | 11.1 | #185 | | |

Peak Table

| Size [nt] | Calibrated Conc. [ng/µl] | Assigned Conc. [ng/µl] | Peak Molarity [nmol/l] | % Integrated Area | Peak Comment | Observations |
|---|---|---|---|---|---|---|
| 25 | 40.0 | 40.0 | 4710 | - | | Lower Marker |
| 1704 | 1.49 | - | 2.57 | 45.30 | | 18S |
| 6383 | 1.80 | - | 0.830 | 54.70 | | 28S |

**Figure S1. RNA quality control and sample preparation. (A)** Ultracentrifuge tubes placed in a snug-fit rack and filled to ~95% of total volume to ensure stability during centrifugation. **(B)** Representative electropherogram displaying RNA size distribution of the human postmortem sample used for polysome profiling.
